# Population-Scale Analysis of Frequency-Dependent Calcium Dynamics in Retinal Ganglion Cells Under Electric Field Stimulation

**DOI:** 10.64898/2026.01.22.701180

**Authors:** Omid Sharafi, Timothy Silliman, Hadi Mokhtari Dowlatabad, Gengle Niu, Steven T. Walston, Jean-Marie C. Bouteiller, Kimberly K. Gokoffski, Gianluca Lazzi

## Abstract

Electric field (EF) stimulation is an emerging neuromodulatory strategy for promoting the repair and functional recovery of degenerated neural networks in neurodegenerative conditions such as glaucoma. EF stimulation therapeutic potential is thought to arise, in part, from modulation of calcium-dependent signaling pathways that regulate neuronal survival and plasticity. However, despite extensive use of EF stimulation in retinal research and clinical studies, it remains unclear how EF waveform frequency and shape govern population-level intracellular calcium dynamics in retinal ganglion cells (RGCs), limiting the rational design of stimulation protocols. Here we address this gap by combining large-scale ex-vivo calcium imaging of Thy1-GCaMP6f mouse retinas with controlled EF stimulation spanning a wide frequency range and a uniform, non-contact stimulation geometry. This approach enables direct measurement of intracellular calcium responses across thousands of individual RGCs under stimulation conditions relevant to non-invasive and translational paradigms. We further develop a morphologically detailed RGC model in NEURON incorporating reaction-diffusion calcium dynamics and admittance-based extracellular stimulation to mechanistically interpret the frequency-dependent responses observed under sinusoidal EF stimulation. Using this integrated experimental–computational framework, we reveal how electric field stimulation modulates population-level calcium signaling in retinal ganglion cells, enabling simultaneous characterization of spatial response patterns and ensemble-averaged activity across thousands of cells. At this scale, RGC calcium responses are constrained to a distinct frequency regime: low frequencies (below 5 Hz) evoke oscillatory transients, intermediate frequencies (10–100 Hz) produce sustained calcium elevation across the population, and high-frequency stimulation (>3 kHz) leads to a sharp attenuation of calcium responses. Among all tested waveforms, a 1:4 asymmetric charge-balanced stimulus at 50 Hz most effectively and consistently elevated intracellular calcium across the RGC population. The computational model reproduces the experimentally observed frequency dependence for sinusoidal stimulation and reveals that the behavior of these different frequency regimes emerges from the interplay between calcium influx, calcium-activated potassium feedback, calcium extrusion kinetics, and soma geometry. Beyond these findings, this work delivers, to our knowledge, the first large-scale dataset of single-cell calcium responses from RGC populations exposed to diverse EF waveforms and frequencies. This dataset enables future data-driven and hybrid modeling approaches that require rich mappings between extracellular stimulation parameters and intracellular calcium dynamics, and establishes a foundation for systematic, physiology-informed optimization of EF stimulation strategies targeting retinal neurodegenerative disease.

## Introduction

Neurons in the central nervous system have a limited capacity to recover once damaged, making neurodegenerative disorders particularly difficult to treat^1^. Glaucoma exemplifies this challenge: long before overt cell loss can be detected, retinal ganglion cells (RGCs) and their circuits begin to undergo functional and structural decline^2,3^. These early circuit-level alterations emerge at stages when no restorative therapies exist, underscoring the need for strategies that can preserve or re-engage neural function. Building on this need for interventions that can act before irreversible cell loss, electrical stimulation of the eye has emerged as a promising approach to modulate RGC activity in glaucoma and related optic neuropathies. Work from our group and others has demonstrated that appropriately structured electric fields can directly influence RGC physiology in ways relevant to repair, with EFs guiding axon growth and promoting directional connectivity in retinal explants^4^, and recent waveform-engineering studies showing that asymmetric charge-balanced stimulation can further enhance these growth-promoting effects^5^. In vivo, extraocular and transcorneal stimulation paradigms slow retinal degeneration and normalize injury-associated epigenetic and transcriptional signatures^6,7^. These findings are consistent with broader preclinical evidence that low-intensity stimulation enhances RGC survival, preserves axons, and reduces inflammatory signaling after optic nerve injury or experimental glaucoma^8–10^. Clinical studies further suggest that alternating-current or transcorneal stimulation improves visual field sensitivity in patients with optic neuropathies and, in some cases, with primary open-angle glaucoma, with larger functional gains in individuals showing stronger stimulation-evoked neural responses^11–13^. Collectively, these findings support the idea that externally applied electric fields can modulate RGC networks in ways that are potentially neuroprotective or restorative.

Mechanistically, electrical stimulation is well positioned to engage intracellular calcium-dependent pathways by modulating membrane potential, recruiting voltage-gated calcium channels, and shaping spatiotemporal calcium fluxes^14,15^. In the retina, prosthetic-like high-frequency and tACS-like electrical stimulation paradigms alter RGC responsiveness and have been linked to calcium-related neuromodulation^16–18^. At the cellular level, many of the pathways implicated in RGC degeneration and survival in glaucoma converge on alterations in intracellular calcium homeostasis. Excessive or dysregulated calcium influx contributes to excitotoxicity and apoptotic cell death in experimental glaucoma and optic nerve crush models^19,20^. Conversely, carefully controlled elevations in calcium can trigger neuroprotective programs that enhance RGC resilience to subsequent injury^19^. Recent work has further shown that variability in RGC homeostatic calcium set points predicts differential vulnerability to degeneration, highlighting calcium as a key determinant of which cells survive under stress^21^. These observations, together with broader evidence that endogenous neuroprotective mechanisms in glaucoma act through calcium-dependent signaling and neurotrophic pathways^22^, suggest that understanding and harnessing calcium dynamics in RGCs is central to designing rational neuroprotective interventions. However, despite growing interest in EF-based therapies for glaucoma and optic neuropathies, it remains poorly understood how the frequency and waveform of EF stimulation govern population-level calcium dynamics in RGCs. Furthermore, we lack a computational model of RGC calcium dynamics that can reveal which biophysical parameters most strongly control these responses and thereby guide the design of optimized waveforms to modulate RGC calcium levels. In this work, we first study how electric field frequency and waveform shape calcium responses in RGC populations by combining large-scale ex-vivo calcium imaging of Thy1-GCaMP6f/+ mouse retinas with an EF stimulation paradigm designed to activate broad RGC ensembles across a wide range of frequencies. The experimental setup employs plate electrodes that are not in direct contact with the retina, making the stimulation geometry more analogous to non-invasive in vivo approaches. This design allows us to characterize how EF waveform frequency transforms RGC calcium responses and how individual cells tile the retina according to their frequency tuning. We then develop a morphologically detailed RGC model in NEURON with RxD-based calcium handling and admittance-derived extracellular stimulation to mechanistically interpret these observations. Once aligned with the experimental recordings, this model provides a platform for systematically probing experimentally untested stimulation scenarios *in silico*, allowing us to predict how calcium dynamics change across waveform and parameter regimes, reduce the number of required animal experiments, and more efficiently converge on candidate stimulation designs for therapy. By probing the roles of calcium and potassium conductances, calcium extrusion kinetics, and soma geometry, the model identifies biophysical parameters that critically shape EF-evoked calcium dynamics. Together, these experimental and modeling results uncover how EF waveform frequency governs RGC calcium activity and identify biophysical parameters that could be targeted to optimize EF-induced engagement of neuroprotective calcium signaling pathways, laying a mechanistic foundation for translating EF stimulation into a rational therapeutic strategy for mitigating glaucomatous neurodegeneration.

## 1 Methods

### 1.1 Experimental Setup

To analyze the frequency characteristics of RGC responses to electrical stimulation, ex-vivo calcium imaging experiments were performed on retinas isolated from transgenic Thy1-GCaMP6f/+ mice (C57BL/6J-Tg(Thy1-GCaMP6f)GP5.17Dkim/J, Jackson Laboratories, Bar Harbour, ME)^23^. Retinas from 10–20 day-old mice were dissected and flattened onto a glass-bottom custom recording chamber. The tissue was oriented with the RGC layer facing downward (toward the glass) to allow imaging from below via an inverted single-photon fluorescence microscope. The retina was held in place at the center of the recording chamber (Fig. 1a), which helped maintain a stable focal plane and high signal-to-noise ratio by minimizing tissue movement. To preserve retinal viability throughout the experiment, circulating Ames’ medium (bubbled with 95% O_2_ and 5% CO_2_) was warmed to 33°C and continuously perfused (5 mL/min) over the tissue. This environment sustained healthy RGC activity for the duration of the recording sessions.

**Figure 1.**
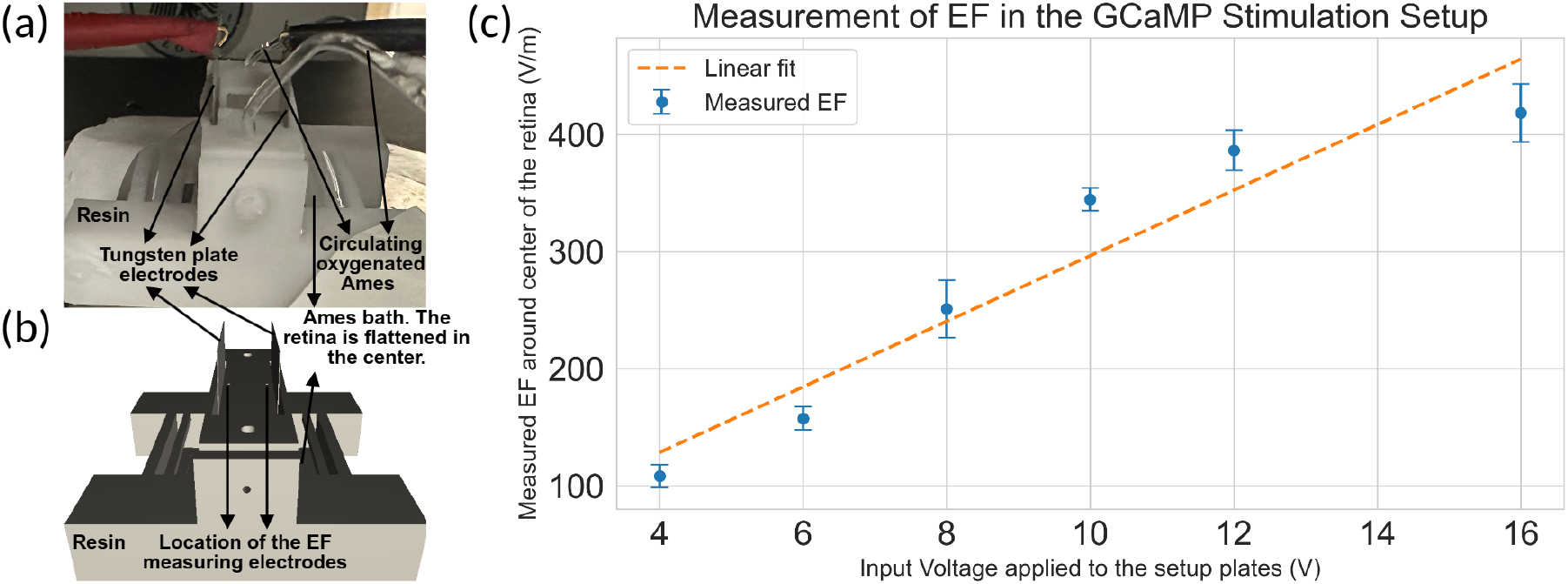
Uniform electric field stimulation of retinal explants. (a) Photograph of the ex-vivo retina experimental setup under the fluorescence microscope. The isolated mouse retina is flattened in the recording chamber with perfusion to maintain viability. (b) Three-dimensional model of the setup used for computational electric field estimation via the admittance method (see Methods). The retina is centered between parallel plate electrodes to create a uniform horizontal field. (c) Measured electric field magnitude at the retina center as a function of applied voltage amplitude, showing 344.61 (±9.85) V/m average induced field for a 10 V sinusoidal stimulus. The dashed red line indicates the least squares linear regression, indicating a linear relationship.

A uniform horizontal EF was applied across the retina using two parallel rectangular 2 cm *×* 2 cm flat tungsten plate electrodes placed 1 cm apart on opposite sides of the chamber. One plate was held at ground potential while stimuli were delivered to the other plate, producing a field oriented tangentially across the retinal layers (orthogonal to the normal photoreceptor-to-RGC signal path). Two measuring electrodes (0.5 mm diameter) were positioned 6.5 mm apart symmetrically around the center of the retina to monitor the local field strength in real time. These measurements confirmed that the induced EF was consistent across different retinas for a given input. To account for any variability in applied EF or optical throughput between preparations, all recorded Thy1-GCaMP6f/+ fluorescence signals were normalized on a per-retina basis before averaging across trials.

The peak voltage of the sinusoidal waveforms was set to 10 V, which generated an average 344.61 (± 9.85) V/m horizontal EF within the tissue, as measured by the probe electrodes. Figure 1a shows a photograph of the experimental setup with stimulation electrodes and perfusion tubing. Figure 1b illustrates a three-dimensional model of the experimental geometry used for computational field calculations, and Fig. 1c shows the measured field strength in the retinal chamber versus input voltage. Least-squares linear regression yielded a coefficient of determination of *R*^2^ = 0.9145, indicating strong linearity.

### 1.2 Electric Field Stimulation Waveform Design

In this study, our stimulation design primarily focused on sinusoidal EFs, as sinusoids provide smoothly varying periodic inputs that enable systematic characterization of frequency-dependent RGC dynamics. Sinusoidal stimulation is widely used in retinal and neural interface studies because its continuous polarity reversal produces predictable membrane polarization, allowing for a direct assessment of how neural populations follow oscillatory fields across various frequencies^24–32^. To capture these dynamics, we selected a set of frequencies spanning low, mid, and high physiological regimes. Low frequencies (1–5 Hz) were chosen to probe slow calcium fluctuations and field-driven modulation on intracellular Ca and GCaMP kinetics timescales. Mid-range frequencies (10–100 Hz) were included based on patch-clamp evidence indicating that most of the RGC types’ maximal spiking rate happens within this band^33^. At the high end, prior work has shown that EFs near 1 kHz can still activate neurons^34^, so we tested 3, 5, and 10 kHz to determine the upper cutoff beyond which RGCs no longer track rapid polarity reversals.

Although sinusoidal stimulation constituted the core of our experimental paradigm, we also included a single asymmetric charge-balanced (ACB) square biphasic pulse as a comparison condition. Phase-asymmetric biphasic pulses have been reported to modulate retinal excitability by shaping cathodic versus anodic polarization^5,30,35^. For this purpose, we implemented a 1:4 ACB waveform at 50 Hz with a -10 V cathodic phase of 2 ms duration, followed by a +2.5 V anodic phase of 8 ms, and a 10 ms rest before the next pulse, making the waveform charge balance and 50% duty cycle, enabling comparison between a sharply polarized biphasic input and the smoothly varying sinusoidal stimuli.

Finally, a short 2-s DC field was applied to evaluate the effect of a non-charge-balanced input relative to the alternating waveforms. The DC duration was intentionally limited to 2 s (shorter than the 10 s duration used for the alternating stimuli) to minimize the risk of sustained polarization-induced cellular damage.

Supplementary Table S1 provides a summary of the stimulation waveform parameters used in our experiments. For sinusoidal waveforms, each cycle included equal-duration anodic and cathodic phases, starting with anodic phase and no rest period, and the number of applied pulses was adjusted according to frequency to maintain the same total induced charge in the 10 s stimulation window. This selection of waveforms allowed us to study the waveform-specific and frequency-specific aspects of calcium dynamics in the RGC population.

### 1.3 Single-Cell Detection and Data Processing

We developed a custom pipeline to detect individual RGCs in the fluorescence images and quantify their calcium response to electrical stimulation. Imaging was performed with an Andor Ixon Ultra EMCCD camera at 512 × 512 pixel resolution using a 20X (NA - 0.75) Nikon Plan Apo objective, which provided a large field of view while resolving single-cell bodies. To aid cell detection, we first computed the temporal standard deviation (S.D.) of fluorescence at each pixel over the course of a recording. This S.D. map highlights active cell locations (which fluctuate in brightness) over the relatively static background. Regions with pronounced intensity variance were fed into a pre-trained U-Net convolutional neural network^36–38^ to segment and identify cell bodies. This automated single-cell detection approach markedly increased the yield and consistency of cell identification across datasets^39^. For each identified cell, raw fluorescence time-series were converted to relative intensity change (Δ*F/F*_0_) traces, which reflect calcium dynamics. Baseline fluorescence *F*_0_ was defined for each cell as the average over 20 frames immediately before EF stimulation onset. The change Δ*F* at any time *t* was then obtained by subtracting this baseline. Each cell’s Δ*F/F*_0_ trace was normalized to its own baseline, and then the responses of all cells in a retina were averaged to yield a population-mean calcium signal for that trial.

Finally, we quantified the total calcium activity induced by a given stimulus using an Averaged Induced Calcium (AIC) metric. AIC is defined as the time-integral of the normalized fluorescence change over the 10 s stimulation period for the population-average trace (Equation 1). AIC measures the aggregate calcium elevation evoked by a stimulus, combining both amplitude and duration of the response into a single value for comparison across conditions.

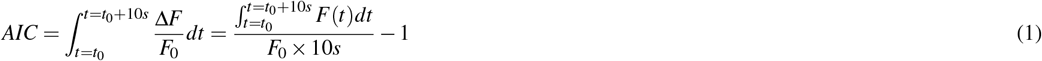

where *t*_0_ is the stimulus onset time. Higher AIC corresponds to a larger overall Ca influx induced in the EF in RGCs. This metric was computed for each stimulus condition in each experiment. The resulting AIC values were averaged across retinas for statistical comparison of waveform efficacy.

### 1.4 RGC Response Stability Protocol and Trial Inclusion Criteria

To assess the stability and viability of retinal ganglion cell (RGC) responses during extended imaging sessions, we periodically delivered a control stimulus using a 1:4 asymmetric charge-balanced (ACB) waveform. This stimulus consisted of a cathodic phase of –10 V, an anodic phase of +2.5 V, and a 50% duty cycle, delivered at 50 Hz. Based on prior work^35^, this waveform reliably evoked calcium transients in RGCs and was used as a benchmark throughout the experiment. These repeated 1:4 control stimuli were delivered at multiple intervals within each session and served as a metric to monitor potential fatigue or response degradation. The consistency of calcium responses to these stimuli confirmed that retinal activity remained stable and that fatigue did not influence activation index (AIC) values.

In addition, sufficient temporal separation was maintained between protocols to avoid interference effects. Each waveform within a protocol was followed by a recovery period, and at least five minutes of rest was given before the next protocol began. This design ensured that neural responses were isolated and unaffected by prior stimulation history. To further ensure data quality, we evaluated the average calcium responses across five repeated control trials within each session. If the response in any trial deviated by more than 20% from the others, the entire session was excluded from subsequent analysis. This quality control step minimized the impact of tissue variability and ensured that only consistent, high-quality data contributed to the final results.

### 1.5 Admittance Method: Constructing the experimental setup and electrodes

A key objective of our modeling framework is to mechanistically interpret how the externally applied electric fields in our ex-vivo preparation translate into membrane polarization within different compartments of an RGC model. To achieve a realistic link between the physical stimulation setup and the compartment-level neural response, we required a modeling framework that could compute the 3D EF distribution generated in Ames’ medium and then inject the resulting currents into a detailed multicompartment neuron model. For this purpose, we used the Admittance Method (AM), a multi-scale biophysical approach that represents the tissue–electrode configuration as a large network of electrical admittances and solves for the voltage at each point in space^40^. Using the AM–NEURON platform^41–44^, the modeling proceeded in two stages. First, the admittance method was used to compute the steady-state voltages and equivalent injected currents induced throughout a 3D volume representing the retina and surrounding medium due to a given stimulus applied at the electrodes. Second, those computed currents were applied as extracellular stimuli in a detailed neuron model of an RGC (described in the next section) to simulate the resulting neural response. For the tissue-level EF calculation, we built a simplified geometry capturing the essential layout of the experimental chamber.

### 1.6 Neuron: Retinal Ganglion Cell Ca model

To investigate the cellular mechanisms underlying the observed calcium signals, we developed a biophysical Ca model of a mouse RGC using the NEURON simulation environment. We used the RGC model established in^45^ using real morphology of RGCs from the NeuroMorpho dataset^46^. D1-bistratified RGCs were modeled to represent average retinal neuronal activity. The model incorporated membrane ion channel dynamics based on Hodgkin–Huxley-style formulations to reproduce RGC electrophysiological behavior^47,48^. In total, seven ionic conductances were included. Five of these were adopted from a standard RGC model defined by Fohlmeister and Miller (providing baseline Na^+^ and K^+^ currents for action potential firing, among others), and two additional currents were added to capture specific RGC response properties. The added currents were a hyperpolarization-activated cation current Ih and a low-voltage-activated T-type calcium current^49–51^.

To simulate calcium transients comparable to those observed in GCaMP6f signals, we incorporated intracellular calcium dynamics into the model. Using NEURON’s Reaction-Diffusion (RxD) module, we created Ca^2+^ handling mechanism: whenever calcium-permeable channels (such as the T-type Ca channel) conducted current, Ca^2+^ was moved from the extracellular space to an intracellular pool in the corresponding compartment. This calcium was then allowed to diffuse and was subject to a decay process representing buffering and pump extrusion. A summary of the mechanisms used in the model is presented in Supplementary Table S2. To translate this intracellular Ca concentration with the recording values, which are proportional to bound GCaMP6f concentration, we used a Hill-type fluorescence model with *K*_*d*_ = 375 nm and Hill coefficient *n* = 2.5 and then calculated AIC based on translated signal^52–54^. Using this framework, we ran simulations for each EF stimulus waveform (using equivalent current injection into the soma calculated from the admittance model as described above) and computed the model-predicted AIC values for comparison with the experimental results.

## 2 Results

### 2.1 AM Confirms Uniform Tangential EF Across Retinal Surface

To demonstrate that our stimulation setup generates a uniform electric field (EF) across the retina, we used the AM to simulate voltage and current flow in the experimental chamber. The model included the electrode layout and the properties of Ames’ medium to calculate how the electric field spreads through the chamber. As shown in Fig. 2a, the voltage map displays a smooth, even gradient between the electrodes, indicating a nearly uniform horizontal EF. Figure 2b shows the current density, with slightly higher currents near the electrodes but mostly uniform flow across the central area where the retina sits, confirming that the RGCs were exposed to a stable, tangential electric field during stimulation. This spatial consistency in the field ensures that the differences in neural responses are due to cell properties rather than an irregular EF.

**Figure 2.**
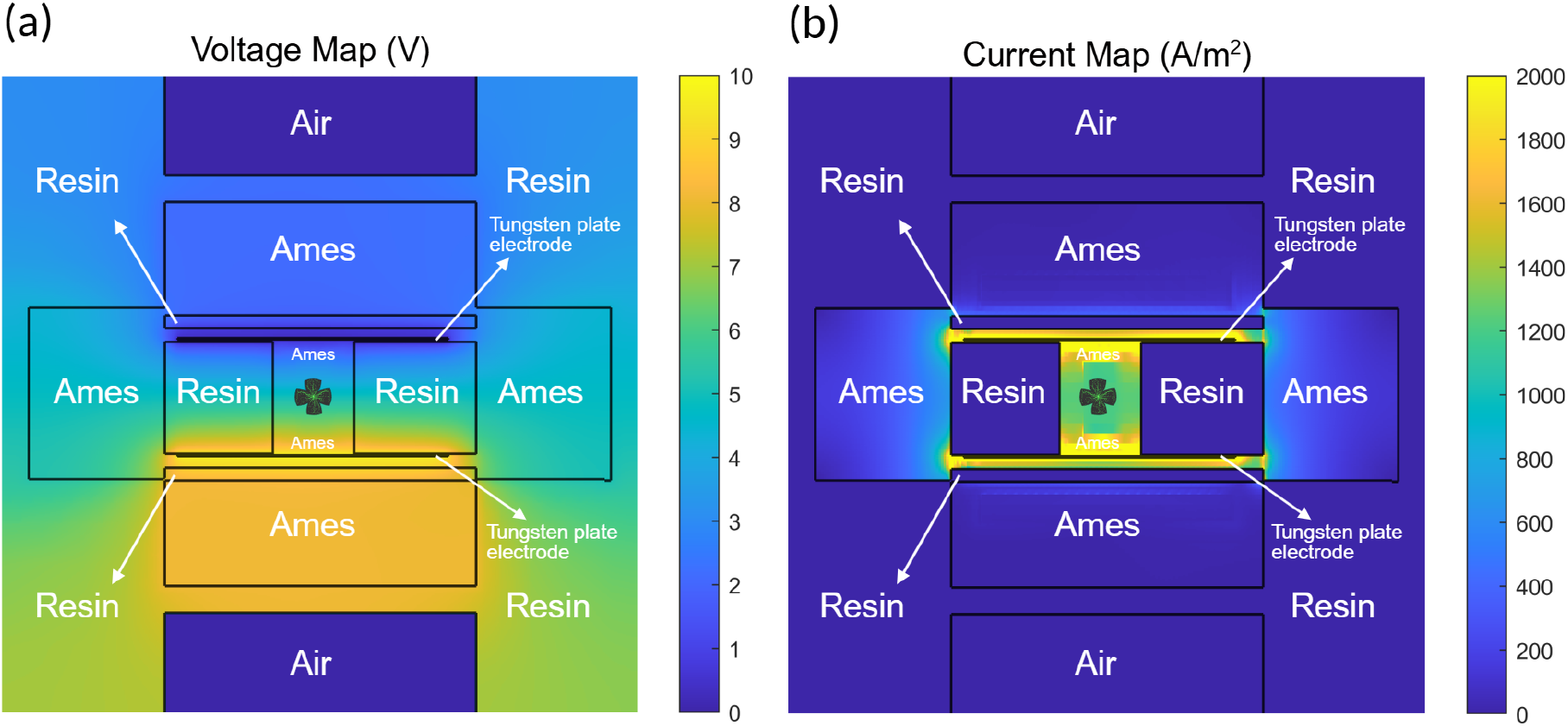
Admittance method simulation of voltage distribution within stimulation chamber. (a) Voltage distribution map across the retinal chamber, showing a near-uniform horizontal field between the electrode plates. (b) Corresponding current density map in *A/m*^2^, indicating regions of elevated current flow near the tissue-electrode interfaces. These maps validate that the stimulation setup produces a well-controlled tangential electric field across the retinal surface.

To evaluate the accuracy of the simulation, we compared the AM-predicted field strength with empirical probe measurements. For a 10 V input, the peak measured field was 640 V/m, while the model predicted 757 V/m, an error of less than 20%. This discrepancy is consistent with expected sources such as electrode-medium interface non-idealities, omission of retina-specific conductivity in the volume model, and minor differences between the true conductivity of Ames’ medium and the literature values used in AM. Overall, the agreement between simulation and measurement indicates that the AM provides a reliable estimate of the field distribution and can therefore be used to link the ex-vivo stimulation paradigm to compartment-level inputs in the RGC computational model.

### 2.2 RGC Responses Remain Stable Over Extended Recordings

To assess whether prolonged electric-field stimulation compromised retinal physiology, we intermittently applied a 1:4 asymmetric charge-balance (ACB) control stimulus at 50 Hz throughout the recording session. This control waveform was chosen because it reliably evokes robust calcium responses without inducing excessive charge accumulation, making it a sensitive indicator of changes in RGC excitability or viability over time.

Across repeated deliveries of the 1:4 ACB control, RGC calcium responses remained stable in both amplitude and temporal profile, with no systematic attenuation or distortion observed as the session progressed (Figure 3). Because these control stimuli were interleaved with higher-frequency sinusoidal and DC test conditions, the preserved response to 1:4 ACB indicates that neither cumulative stimulation nor extended imaging led to measurable degradation of RGC physiological responsiveness. Consistent stability was observed at both the single-cell and population levels, and trials exhibiting irregular or weakened responses were excluded from subsequent analyses. Together, these results confirm that the retina remained functionally healthy throughout the experiment and that the reported effects of test stimuli are unlikely to arise from stimulation-induced toxicity or recording-related drift.

**Figure 3.**
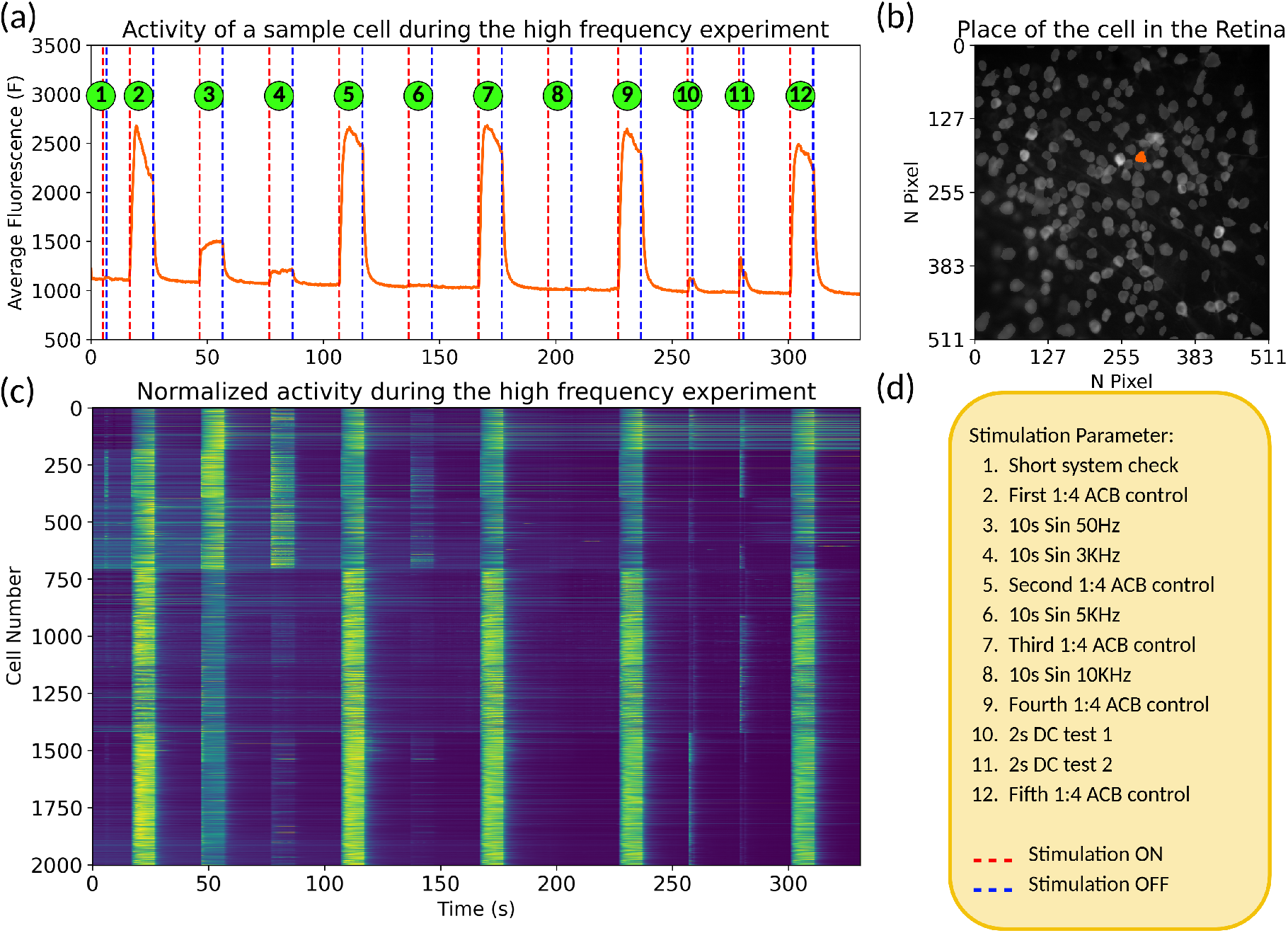
RGC responses are stable over repeated 1:4 ACB stimulations. (a) Average raw fluorescence trace (f(t)) from a representative RGC exposed to high-frequency sinusoidal stimulation. 1:4 ACB at 50 Hz was delivered intermittently to confirm RGC health. The consistency in amplitude and shape of the calcium transients across these repeated 1:4 ACB controls indicates that the retina maintained stable physiological responsiveness throughout the recording session. (b) Spatial location of the representative RGC within the imaged retinal field. The orange marker indicates the centroid of the cell whose fluorescence trace is shown in panel (a). (c) Heatmap showing normalized f(t) across all detected RGCs during the same examination protocol. Trials with inconsistent or diminished population responses were excluded from subsequent analyses to ensure that only robust, high-quality recordings contributed to the reported results. (d) Summary of the stimulation protocol used in this experiment. The session consisted of intermittent 1:4 ACB control pulses delivered between sinusoidal and DC test stimuli to verify retinal health throughout the recording. Red dashed lines denote stimulation onset, and blue dashed lines denote stimulation offset.

### 2.3 Stimulation Frequency Modulates Both Strength and Recruitment Diversity of Individual RGC Responses

To understand how RGCs respond to different frequencies of EF stimulation, we analyzed calcium activity at the single cell level. While the average population response shows general trends, looking at single-cell responses helps reveal how much variation exists between RGCs and which stimulation patterns are more broadly effective. Figure 4(a) shows violin plots of the average integrated Δ*F/F*_0_ (AIC) responses from individual RGCs for each sinusoidal frequency. At low frequencies (1–5 Hz), most cells showed small responses with values tightly grouped around the median, suggesting weaker and more uniform activation. In the mid-frequency range (10–100 Hz), the responses were stronger and more spread out, indicating that more cells were activated and with a wider range of response levels. At high frequencies (≥ 3 kHz), the responses became small and tightly clustered again, showing that very few cells could follow the fast stimulation.

**Figure 4.**
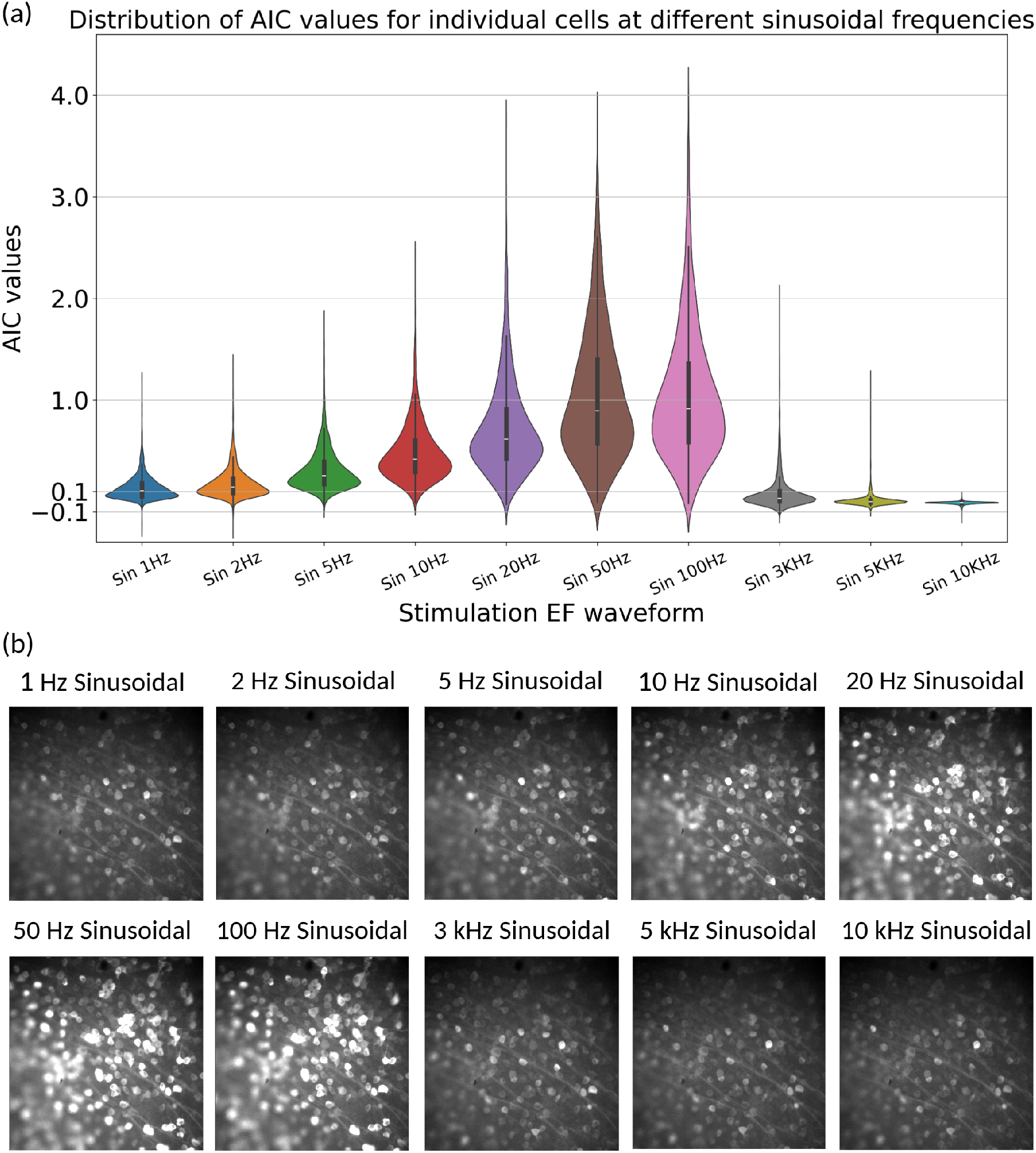
Population response of RGCs to sinusoidal stimulation. (a) Single-cell calcium response varies across sinusoidal stimulation frequencies. Violin plots illustrate the distribution of average integrated Δ*F/F*_0_ responses from individual RGCs in a representative retina subjected to sinusoidal EF stimulation at varying frequencies (1 Hz to 10 kHz). The box values represent the first quartile, median, and third quartile. The distribution shapes at lower frequencies (1–10 Hz) and at high-frequency stimulations (≥ 3 kHz) are more concentrated around the median, while mid-range frequencies (20–100 Hz) result in a stronger and more spread-out distribution of cell activity. Additionally, in all frequencies, some cells exhibit high activity, while others remain inactive, confirming the variability in activity among different RGC cell types. (b) A sample raw recording section of peak values for one trial at different frequencies is presented.

Across all frequencies, we also observed that some cells showed strong activity while others remained inactive. Given that cells were exposed to a uniform EF, this pattern highlights the natural diversity among RGCs; some cell types or positions in the retina may be more responsive to electric stimulation than others. These results emphasize the importance of choosing stimulation parameters that activate a broad range of RGCs, not just a small subset. Figure 4(b) provides an example of a raw recording, highlighting the peak response values across different frequencies. This example illustrates how much calcium influx can vary between cells. Overall, these findings confirm that the frequency of stimulation controls not just how much activity occurs, but also how many and which RGCs are recruited.

### 2.4 RGCs Preferentially Respond to Distinct EF Frequencies and do not cluster within the Retina

To further investigate RGC diversity in response to EF stimulation, we analyzed the spectrum of RGC responses across the retina. In Figure 5(a), we mapped the normalized activity of individual RGCs in response to a 100 Hz sine wave stimulus across multiple retinas. To better visualize the variation, calcium responses (AIC values) were quantile-transformed into 40 equally populated bins and then linearly scaled. This normalization method reveals the relative distribution of activity and shows that response strength varies smoothly across the retinal surface, suggesting a distributed organization of EF sensitivity among RGCs. This spatial tiling is suggestive of the functional mosaics observed in light-evoked responses of RGC subtypes^55^.

**Figure 5.**
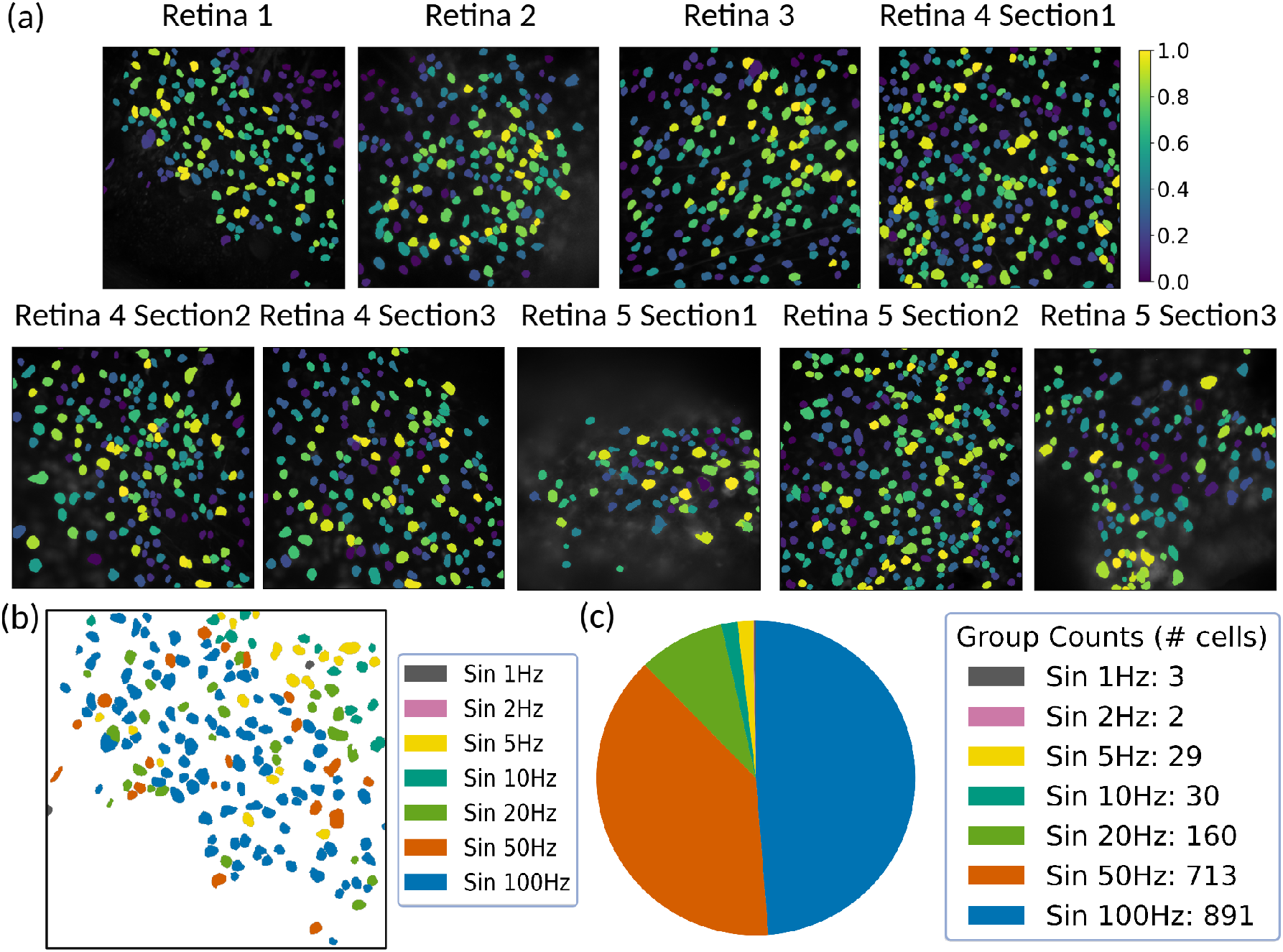
Spatial distribution of RGC response gradients. (a) Spatial maps of individual RGC responses to a 100 Hz sine wave stimulus across sections from five retinas. AIC values were quantile-normalized (40 bins) to highlight response gradients. Warmer colors indicate higher normalized activity. (b) Color-coded map showing each RGC labeled by the frequency at which it exhibited its peak AIC response. Cells preferring different frequencies tile the retinal space, illustrating functional diversity. (c) Pie chart showing the number of cells whose peak response occurred at each stimulus frequency. The majority of RGCs showed maximal activation at 50 or 100 Hz, though a smaller population was tuned to lower frequencies.

To explore which EF frequency each cell responded to most strongly, we identified the frequency that evoked the maximum AIC value for each RGC. Figure 5(b) color-codes individual cells according to this frequency preference. Most cells showed peak responses at either 50 Hz or 100 Hz, but a subset of cells responded more strongly to lower frequencies, indicating functional diversity. Notably, cells preferring different frequencies were spatially intermingled, suggesting that RGC subtypes with distinct frequency tuning tile the retina in a mosaic-like fashion. Figure 5(c) quantifies the frequency preferences across the dataset. Of 1,828 analyzed cells, 87.7% showed peak responses at 50 or 100 Hz, consistent with the high efficacy of these frequencies in population-level analysis. However, the remaining cells peaked at lower frequencies (1–20 Hz), supporting the idea that different RGC types are selectively tuned to distinct EF frequency ranges.

### 2.5 RGCs Preferentially Respond to Distinct EF Frequencies and do not cluster within the Retina

To further investigate RGC diversity in response to EF stimulation, we analyzed the spectrum of RGC responses across the retina. In Fig. 5(a), we mapped the normalized activity of individual RGCs in response to a 100 Hz sine wave stimulus across multiple retinas. To better visualize the variation, calcium responses (AIC values) were quantile-transformed into 40 equally populated bins and then linearly scaled. This normalization method reveals the relative distribution of activity and shows that response strength varies smoothly across the retinal surface, suggesting a distributed organization of EF sensitivity among RGCs. This spatial tiling is suggestive of the functional mosaics observed in light-evoked responses of RGC subtypes^55^.

To explore which EF frequency each cell responded to most strongly, we identified the frequency that evoked the maximum AIC value for each RGC. Figure 5(b) color-codes individual cells according to this frequency preference. Most cells showed peak responses at either 50 Hz or 100 Hz, but a subset of cells responded more strongly to lower frequencies, indicating functional diversity. Notably, cells preferring different frequencies were spatially intermingled, suggesting that RGC subtypes with distinct frequency tuning tile the retina in a mosaic-like fashion. Figure 5(c) quantifies the frequency preferences across the dataset. Of 1,828 analyzed cells, 87.7% showed peak responses at 50 or 100 Hz, consistent with the high efficacy of these frequencies in population-level analysis. However, the remaining cells peaked at lower frequencies (1–20 Hz), supporting the idea that different RGC types are selectively tuned to distinct EF frequency ranges.

### 2.6 RGC Calcium Dynamics Reflect Stimulus Waveform, with Intermediate-Frequency Sinusoids Inducing Sustained Elevation

Having established stable recording conditions, we next examined how RGC calcium signals depended on the shape and frequency of the EF stimulus. We applied a variety of horizontal EF waveforms to the same retina and observed distinctly different calcium response profiles for each stimulus condition (Figure 6). Notably, the temporal pattern of the Ca signal closely reflected the input waveform in many cases. For low-frequency sinusoidal stimulation (1–5 Hz), the RGCs exhibited oscillatory calcium responses that rose and fell in phase with each cycle of the field. These slow oscillations allowed Ca levels to return near baseline between stimuli, producing a clear periodic modulation in the Δ*F/F*_0_ trace.

**Figure 6.**
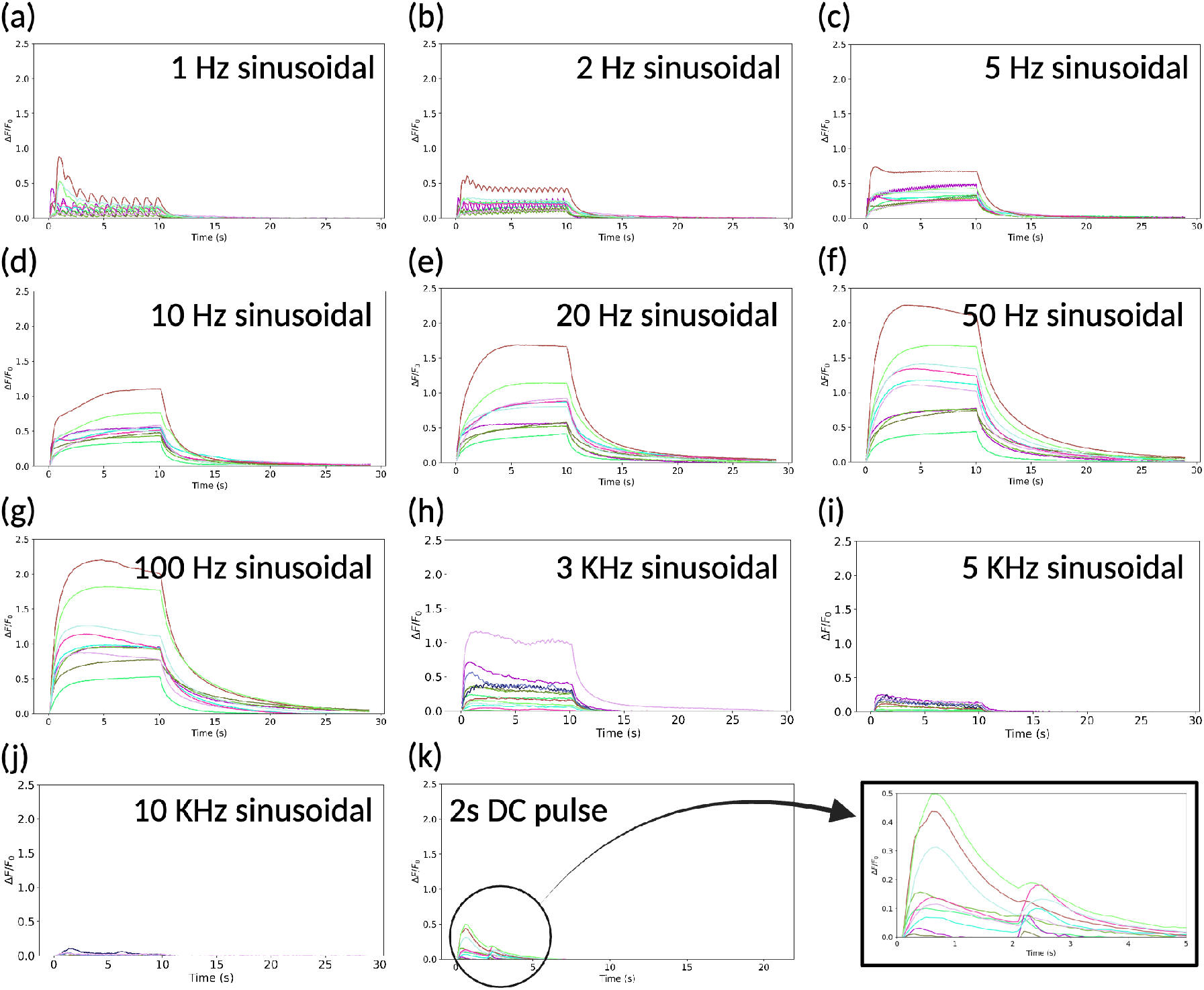
RGC calcium signal time-courses under different EF stimulation waveforms. Each colored trace represents the average normalized Δ*F/F*_0_ response from one recording session (over 200 cells each session) to a given waveform. Stimuli tested include: (a) 1 Hz sinusoidal, (b) 2 Hz sinusoidal, (c) 5 Hz sinusoidal, (d) 10 Hz sinusoidal, (e) 20 Hz sinusoidal, (f) 50 Hz sinusoidal, (g) 100 Hz sinusoidal, (h) 3 kHz sinusoidal, (i) 5 kHz sinusoidal, (j) 10 kHz sinusoidal, and (k) a 2 s DC pulse. The temporal profiles of the Ca responses differ markedly between low-frequency (periodic spiking and full relaxations), mid-frequency (sustained elevation with partial modulation), high-frequency (diminished response), and DC (transient adapting response) stimulations.

At intermediate frequencies (10–100 Hz sinusoids), the Ca traces showed a fused but still robust elevation – the cells could not fully relax between pulses, leading to a summation of Ca transients and a higher trace during stimulation. For example, a 50 Hz sinusoidal field induced a sustained calcium elevation with superimposed small ripples corresponding to the stimulus cycles. Extremely high-frequency stimulation, however, failed to maintain the calcium response. When 3 kHz and 5 kHz sinusoidal fields were applied, the RGCs still showed an initial rise in Ca^2+^, but the response was at a much lower level and began to vanish despite continued stimulation. By 10 kHz, the cells displayed little to no calcium increase above baseline – effectively, the RGCs could not follow such rapid electric field oscillations, leading to a negligible net calcium influx.

Finally, the 2-second DC step (a constant electric field) produced a transient calcium response characterized by an initial sharp rise followed by adaptation: the Ca signal peaked quickly and then partially decayed during the 2-second interval. Importantly, the qualitative shape of each Ca response (oscillatory vs. sustained vs. adapting) was reproducible across retinas, even though absolute response magnitudes varied with factors like baseline fluorescence intensity and cell yield. In all retinas tested, we observed the same rank-order of responsiveness for the different waveforms, indicating that the effects of stimulus waveform on RGC calcium dynamics are robust and generalizable.

### 2.7 AIC Analysis Reveals Optimal Frequency and Waveform for Retinal Ganglion Cell Activation

To quantitatively compare the efficacy of each stimulation pattern, we computed the averaged induced calcium (AIC) for each waveform across all experiments. Over 2000 individual RGCs were detected and analyzed. All AIC values were normalized to the 50 Hz sinusoidal baseline (yellow bar in Fig. 7) for ease of comparison. We found that the stimulus waveform had a profound impact on the total Ca^2+^ influx (Figure 7). The 1:4 ACB pulse train at 50 Hz produced the highest AIC of all conditions, on average 130% (± 58%) greater than that of the 50 Hz sinusoid (baseline). This indicates that the asymmetric biphasic pulses were significantly more effective at recruiting calcium responses in RGCs than a simple harmonic field at the same frequency.

**Figure 7.**
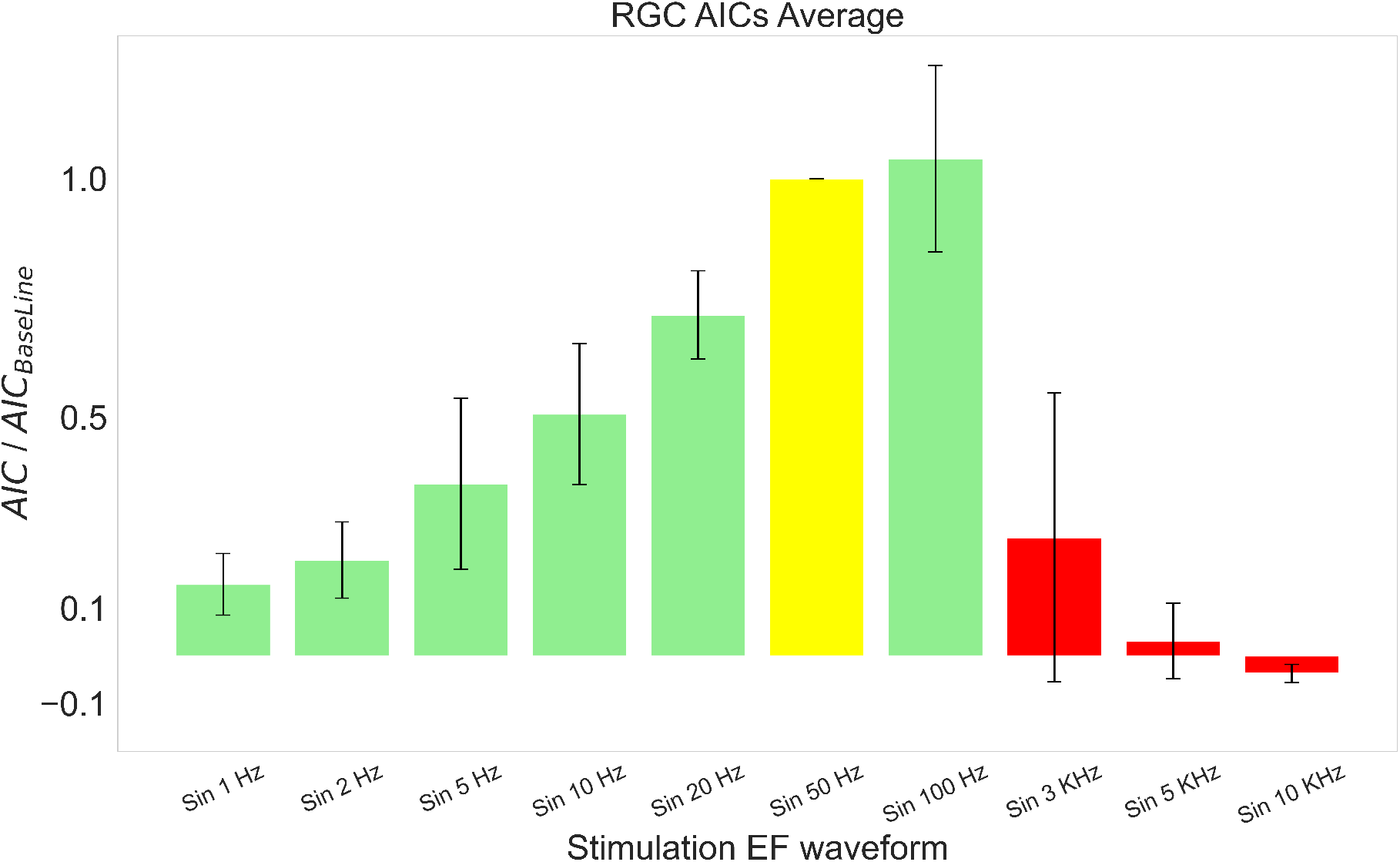
Quantitative comparison of RGC calcium responses (AIC metric) across stimulation frequencies. Values are normalized to the 50 Hz sinusoidal case (yellow bar = 1.0). Average AIC values for high-frequency and ACB waveforms. Red bars show AIC for high-frequency sinusoids (3 kHz, 5 kHz, 10 kHz), all of which are much lower – the 3 kHz sinusoid produces only 0.2 relative AIC, the 5 kHz nearly zero, and negative activity for the 10 kHz defining a cutoff frequency around 3 kHz. AIC values for low-frequency sinusoidal stimulation (1, 2, 5, 10, 20 Hz; green bars). These responses are also weaker than the 50 Hz baseline (e.g., 10 Hz yields 0.5, and 1 Hz 0.1), reflecting insufficient temporal summation at low rates. Error bars indicate ±1 SD across n = 5 retinas per condition.

Examining the sinusoidal frequency sweep, we identified an optimal range around tens of Hz for calcium accumulation. The 50 Hz sinusoid elicited one of the largest Ca responses among the sine waves tested. When frequency was reduced below 50 Hz, the AIC dropped progressively – e.g., 20 Hz and 10 Hz yielded only about 70% and 50%, respectively, of the 50 Hz reference AIC. At the lowest frequencies (1–5 Hz), AIC values were minimal (10–20% of baseline), since the cells had ample time to recover between stimuli and thus produced only transient Ca spikes with little summation.

On the other extreme, pushing the frequency higher also caused a decline in effectiveness. Notably, the cutoff frequency for sinusoidal stimulation was around 3 kHz: stimuli at 3 kHz still induced a measurable Ca elevation (though only 20% of the 50 Hz reference), but at 5 or 10 kHz the AIC was nearly zero (no significant Ca increase). This sharp drop-off at high frequencies likely corresponds to the inability of RGC membranes to respond to stimuli faster than their membrane time constants^33^. In summary, the AIC analysis indicates that a mid-range frequency (approximately 50 Hz) with an asymmetric waveform is the most efficient for driving calcium entry in RGCs under our experimental conditions. The 1:4 ACB at 50 Hz was the top performer, significantly outperforming both lower-frequency stimuli (which lack temporal summation) and higher-frequency stimuli (which exceed the cells’ following capability). These findings quantitatively substantiate the notion of waveform-specific calcium dynamics, demonstrating that careful selection of stimulus parameters can drastically influence the level of neural activation achieved.

### 2.8 Computational Prediction of Frequency-Dependent Calcium Dynamics in RGCs

To elucidate which ion channels contributed to our experimental findings, we used a biophysical computational model of a D1-type RGC implemented in the NEURON simulator with the RxD module for calcium dynamics. D1-type RGCs are well-characterized bistratified cells with relatively large somas and prominent dendritic trees, making them suitable for modeling stimulus-evoked calcium accumulation under extracellular EF stimulation^56^. This model incorporates realistic ion channel kinetics and morphology, allowing us to simulate intracellular calcium accumulation in response to EF stimulation. We subjected the model to sinusoidal EF waveforms across a broad range of frequencies (1 Hz to 10 kHz) and computed the predicted calcium activity using the same Averaged Induced Calcium (AIC) metric used in our experiments. All AIC values were normalized to the value obtained for the 50 Hz sinusoidal waveform, which served as the baseline condition.

As shown in Fig. 8(a), the model replicated key trends observed experimentally. AIC increased with frequency up to 100–200 Hz, peaking slightly above the 50 Hz baseline, then declined at higher frequencies. The model showed a steep drop-off in calcium activity beyond 1–2 kHz, with virtually small negative response at 5 and 10 kHz, consistent with the high-frequency cutoff observed in ex-vivo recordings. The model predicts the cut-off frequency to be around 2 kHz, which is lower than the experimental values of 3 kHz. This discrepancy can be explained by the fact that we are using only one type of RGC while the recording contains different types of responses, as illustrated in Fig. 4.

**Figure 8.**
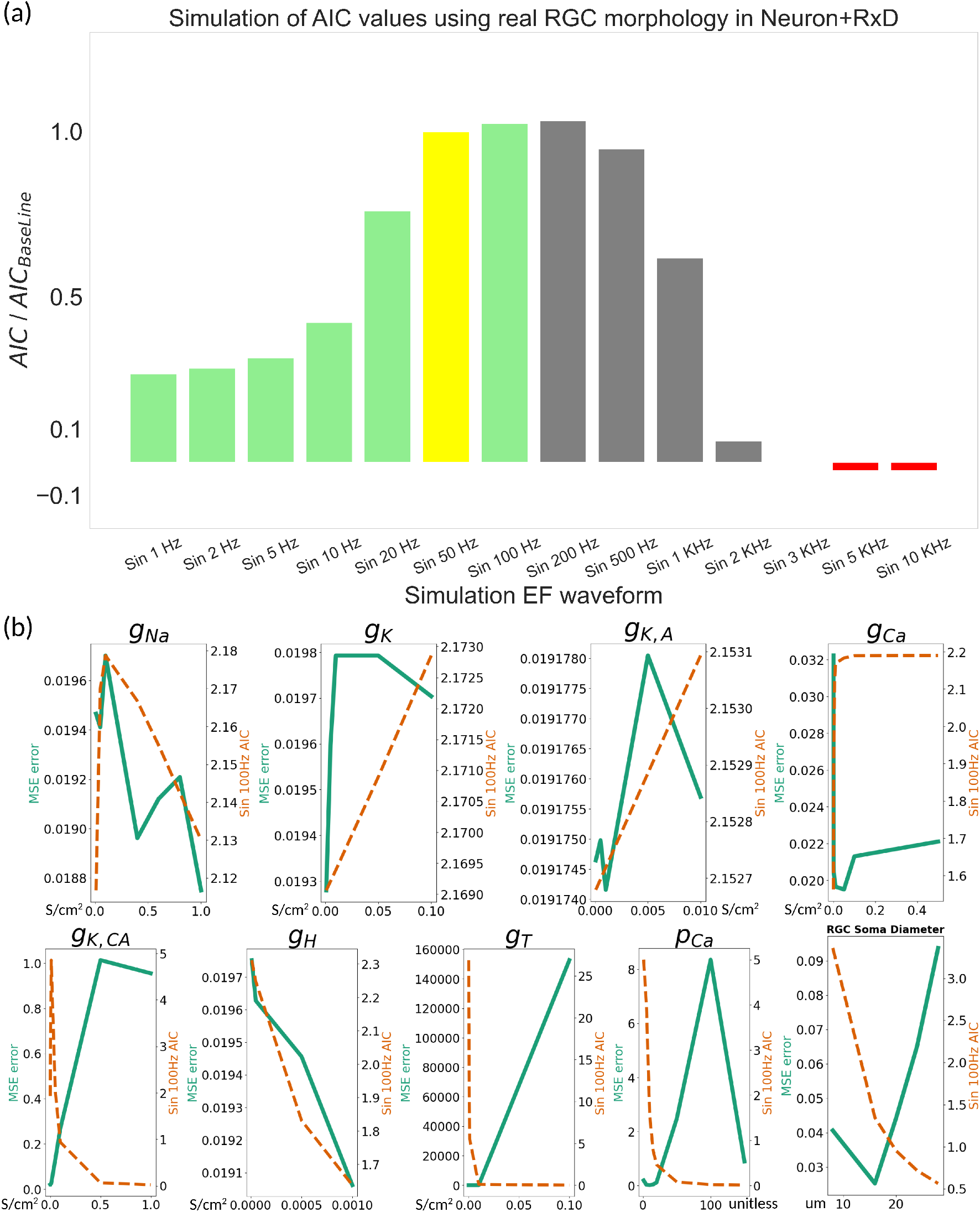
Simulation of RGC calcium dynamics using a biophysical model. (a) Simulated AIC values in response to sinusoidal EF waveforms (1 Hz to 10 kHz) normalized to the 50 Hz response (yellow bar). Bars in gray represent frequencies not tested experimentally. The model predicts frequency tuning consistent with experimental findings: peak responses around 50–200 Hz and sharp decline beyond 2 kHz. (b) Sensitivity analysis of key biophysical parameters. For each parameter, the effect on model MSE error (green) and the predicted AIC at 100 Hz (orange dashed line) is shown. Parameters like Ca-activated potassium channel conductance, high-voltage-activated Calcium channel conductance, T-type Calcium channel conductance, Ca extrusion pump, and A-type potassium conductance, as well as soma size, significantly modulate calcium responses.

To further explore how specific biophysical properties contribute to the frequency tuning of calcium signals, we performed a parameter sensitivity sweep across nine variables: *g*_Ca_, *g*_K,Ca_, *g*_Na_, *g*_K_, *g*_T_, *g*_H_, calcium pump rate (*p*_Ca_), and soma diameter (Fig 8(b)). For each parameter, we quantified (1) the model-predicted AIC at 100 Hz (orange dashed line) and (2) the mean squared error (MSE) between the model-predicted AIC frequency profile and the experimental profile (green line). The sweep revealed several distinct patterns. Increasing *g*_Ca_ boosted the calcium response at 100 Hz and reduced MSE up to an optimal value, consistent with stronger calcium entry improving agreement with the data, but responses became large when *g*_Ca_ was too high so the MSE error started to increase again. Increasing *g*_K,Ca_ in a very small window at the beginning of the sweep showed a rapid rise in calcium response, but after that, the MSE error increased and the calcium response decreased monotonically. This suggests that there is narrow range of physiologically meaningful conductance for *g*_K,Ca_ and reflects the suppressive action of calcium-activated potassium currents on calcium buildup. *g*_T_ showed the same trend but in a more extreme scenario. As *g*_T_ moves out of the sweet range, the calcium trace converges to zero and the MSE error increases rapidly. This effect can be due to the initial large T-type calcium influx that strongly activates calcium-dependent potassium channels (*g*_K,Ca_). If that potassium feedback is strong, the cell is pulled down to more negative voltages, making it even harder for calcium channels (both T-type and high-voltage ones) to reopen during subsequent cycles. *g*_Na_, *g*_K_, and *g*_K,A_ produced only small shifts in MSE, suggesting they contribute indirectly through excitability rather than directly shaping calcium accumulation. *g*_H_ also had limited influence, mainly shifting baseline excitability without altering the frequency profile. Both the calcium pump and soma diameter had significant effects on AIC and MSE errors. Higher *p*_Ca_, which means more powerful ATP pump calcium extrusion, reduced the calcium signal at 100 Hz monotonically and caused a peak in MSE error, showing that fast removal of calcium prevents the buildup seen in experiments. Increasing soma diameter decreased the overall amplitude of the response monotonically. This is completely aligned with previous studies^57,58^. MSE error was higher than 0.4 when diameter is less than 10 *µ*m or above 20 *µ*m. This range is compatible with mouse RGC sizes, as most of the RGC types have soma sizes within in this range.

## 3 Discussion

Development of successful electrical stimulation approaches for neuro-degenerative diseases requires establishing stimulation conditions that are wide reaching. Neural cellular diversity complicates this goal as one stimulation approach is unlikely to be effective for all cell types and conditions. This study provides a comprehensive investigation into how EF waveforms and frequency affect calcium signaling in RGCs. By integrating ex-vivo calcium imaging with high spatial resolution and NEURON-based computational modeling, we identified frequency-dependent calcium dynamics that are strongly shaped by both the waveform and stimulation rate. Our findings show that a 1:4 asymmetric charge-balanced (ACB) pulse train at 50 Hz elicits the strongest and most consistent calcium responses among the tested conditions. In contrast, sinusoidal waveforms reveal a clear frequency tuning, with peak responses in the 20–100 Hz range and a sharp attenuation beyond 3 kHz.

These insights have direct implications for improving the effectiveness of EF stimulation as a treatment for neurodegenerative diseases like glaucoma. By selecting waveform shapes and frequencies that maximize RGC activation within physiological constraints, electrical stimulation strategies can play a critical role in restoring degenerating neural networks. Our work emphasizes that the same total charge delivered in different temporal patterns can result in dramatically different neural outcomes.

To better understand the role that different ionic channels play in mediating cellular responses to EF stimulation, we developed a generalizable computational approach using NEURON’s RxD framework. The model replicated a peak in calcium response between 50–200 Hz and a sharp drop beyond 3 kHz. A parameter sweep revealed that membrane conductances, especially those involving calcium and potassium channels, and soma morphology play a critical role in shaping these dynamics.

These results are significant as they highlight how the interplay between calcium entry and potassium-mediated feedback sets the operating range of RGC responses. Calcium conductances (*g*_Ca_, *g*_T_) provide the primary influx needed to generate robust signals, but they also recruit calcium-dependent potassium currents (*g*_K,Ca_) that impose strong negative feedback^56^. This balance amplifies responses within the physiological frequency range (50–200 Hz) while preventing excessive accumulation at higher intensities. Once stimulation exceeds this regime, the feedback dominates, hyperpolarizing the membrane and silencing subsequent channel activity, which explains the sharp collapse of calcium responses beyond 2 kHz. Soma morphology further modulates this balance, with larger diameters diluting intracellular calcium and reducing response amplitude, whereas smaller diameters promote stronger but less stable accumulations. The optimal range of 12–20 *µ*m closely matches the physiological soma sizes of most mouse RGCs.

Although this study focused on D1-type RGCs, the modeling framework is broadly applicable and can be extended to a wide range of neuronal cell types and stimulation paradigms, including other regions of the central nervous system. This versatility makes the approach a valuable tool for probing activity-dependent calcium dynamics across diverse neuronal populations. Our computational results not only mimic our experimental observations but also provide mechanistic insights into how different waveform parameters interact with the cell’s biophysical and morphological properties to influence RGC excitability and calcium dynamics. Overall, these results indicate that the model constitutes a predictive tool that may be used to guide the design of future stimulation protocols and predict how the application of an EF with the same total charge, but delivered with different temporal profiles, can result in dramatically different neural outcomes. The computational insights have direct applications for optimizing EF-based strategies for neuro-restoration by aligning stimulation strategies with the natural biophysical tuning of retinal neurons.

Future work will expand this approach in several directions. First, we plan to simulate and experimentally validate additional waveform families, including other ACB configurations and short biphasic pulses. Second, future experiments will examine cell-type-specific responses by leveraging morphological and light response data, allowing for more tailored stimulation strategies. Third, we will explore the interaction of spatial and temporal stimulation patterns in multi-electrode or patterned EF delivery, aiming to further improve focality and efficiency. Finally, we intend to integrate closed-loop modeling and simulation in real time, enabling adaptive waveform optimization based on observed calcium feedback. On the modeling end, by advancing the mechanisms used in the computational model and incorporating additional morphological RGC types, we aim to improve its generality and predictive accuracy for calcium response dynamics across the broader RGC population. Together, these efforts aim to establish a generalizable platform for data-driven, physiologically informed design of neural stimulation protocols for vision restoration.

## Acknowledgements

KKG and GL were supported by grants from NEI/NIH (R01EY035375) and NSF (2121164). KKG was also supported by a grant from the Research to Prevent Blindness Foundation Disney Award for Amblyopia. OS was supported by a grant from the NSF (2121164). This work was also supported by an unrestricted grant to the Department of Ophthalmology from Research to Prevent Blindness and the NEI (P30EY029220). The content is solely the responsibility of the authors and does not necessarily represent the official views of the National Institutes of Health. No funding sponsors were involved in the study design, collection, analysis, interpretation of data, writing of the report, or decision to submit the article for publication.

## Author contributions statement

O.S. conceived and designed the experiments. T.S. prepared the experimental setup. G.N. prepared retinal tissue and performed retina dissection and flattening. O.S. conducted the experiments, analyzed the data, developed the computational model, and prepared the manuscript and figures. H.M. contributed to electrode measurements and electric field simulations. S.W. and K.G. provided experimental guidance and supervision. J.B. advised on the computational modeling. G.L. provided overall supervision and critical feedback on the study. All authors reviewed and approved the final manuscript.

## Additional information

### Data and code availability

Raw calcium imaging recordings (`.tiff`), representative supplementary videos (`.mp4`), processed single-cell time series (`Python pickle` format), and all data-processing pipelines (`Python`) will be provided publicly after acceptance in the journal. The computational materials, including the morphologically detailed `NEURON+RxD` calcium model, the admittance-method–based electric field calculation code, and the notebooks used to generate the figures, are also included. All data and code will be made publicly available on GitHub upon acceptance of the manuscript.

## References

1. He, Z. & Jin, Y. Intrinsic control of axon regeneration. Neuron 90, 437–451, DOI: 10.1016/j.neuron.2016.04.022 (2016).

2. Crish, S. D. & Calkins, D. J. Central visual pathways in glaucoma: Evidence for distal mechanisms of neuronal self-repair. J. Neuro-Ophthalmology 35, S29–S37, DOI: 10.1097/wno.0000000000000291 (2015).

3. Buckingham, B. P. et al. Progressive ganglion cell degeneration precedes neuronal loss in a mouse model of glaucoma. J. Neurosci. 28, 2735–2744, DOI: 10.1523/JNEUROSCI.4443-07.2008 (2008).

4. Gokoffski, K. K., Jia, X., Shvarts, D., Xia, G. & Zhao, M. Physiologic electrical fields direct retinal ganglion cell axon growth in vitro. Investig. Ophthalmol. & Vis. Sci. 60, 3659–3668, DOI: 10.1167/iovs.18-25118 (2019).

5. Peng, M. G. et al. Asymmetric charge balanced waveforms direct retinal ganglion cell axon growth. Sci. Reports 13, 13233, DOI: 10.1038/s41598-023-40097-6 (2023).

6. Gonzalez Calle, A. et al. An extraocular electrical stimulation approach to slow down the progression of retinal degeneration in an animal model, DOI: 10.1038/s41598-023-40547-1 (2023).

7. Tew, B. Y. et al. Transcorneal electrical stimulation restores DNA methylation changes in retinal degeneration. Front. Mol. Neurosci. 17, 1484964, DOI: 10.3389/fnmol.2024.1484964 (2024).

8. Morimoto, T. et al. Transcorneal electrical stimulation rescues axotomized retinal ganglion cells by activating endogenous retinal IGF-1 system. Investig. Ophthalmol. & Vis. Sci. 46, 2147–2155, DOI: 10.1167/iovs.04-1339 (2005).

9. Yin, H. et al. Transcorneal electrical stimulation promotes survival of retinal ganglion cells after optic nerve transection in rats accompanied by reduced microglial activation and TNF-α expression. Brain Res. 1650, 10–20, DOI: 10.1016/j.brainres.2016.08.034 (2016).

10. Jassim, A. H., Cavanaugh, M., Shah, J. S., Willits, R. K. & Inman, D. M. Transcorneal electrical stimulation reduces neurodegenerative process in a mouse model of glaucoma. Annals Biomed. Eng. 49, 858–870, DOI: 10.1007/s10439-020-02608-8 (2021).

11. Gall, C. et al. Alternating current stimulation for vision restoration after optic nerve damage: a randomized clinical trial. PloS one 11, e0156134, DOI: 10.1371/journal.pone.0156134.

12. Schmidt, S. et al. Progressive enhancement of alpha activity and visual function in patients with optic neuropathy: a two-week repeated session alternating current stimulation study. Brain Stimul. 6, 87–93, DOI: 10.1016/j.brs.2012.03.008 (2013).

13. Ota, Y. et al. The efficacy of transcorneal electrical stimulation for the treatment of primary open-angle glaucoma: A pilot study. The Keio J. Medicine 67, 45–53, DOI: 10.2302/kjm.2017-0015-OA (2017).

14. Kadan-Jamal, K. et al. Electrical stimulation of cells: Drivers, technology, and effects. Chem. Rev. 125, 6874–6905, DOI: 10.1021/acs.chemrev.4c00468 (2025).

15. Ullrich, V. & Apell, H.-J. Electromagnetic fields and calcium signaling by the voltage dependent anion channel. Open J. Vet. Medicine 11, 57–86, DOI: 10.4236/ojvm.2021.111004 (2021).

16. Peiroten, L., Zrenner, E. et al. Artificial vision: The high-frequency electrical stimulation of the blind mouse retina—decay, spike generation, and electrogenically clamped intracellular Ca2+ at elevated levels. Bioengineering 10, 1208, DOI: 10.3390/bioengineering10101208 (2023).

17. Amthor, F. R. & Strang, C. E. Effects of tacs-like electrical stimulation on on-center retinal ganglion cells: Part i. Eye Brain 13, 175–192, DOI: 10.2147/EB.S312402 (2021).

18. Weitz, A. C. et al. Imaging the response of the retina to electrical stimulation with genetically encoded calcium indicators. J. Neurophysiol. 109, 1979–1988, DOI: 10.1152/jn.00852.2012 (2013).

19. Brandt, S. K., Weatherly, M. E., Ware, L., Linn, D. M. & Linn, C. L. Calcium preconditioning triggers neuroprotection in retinal ganglion cells. Neuroscience 172, 387–397, DOI: 10.1016/j.neuroscience.2010.10.071 (2011).

20. Quigley, H. A. Neuronal death in glaucoma. Prog. Retin. Eye Res. 18, 39–57, DOI: 10.1016/S1350-9462(98)00014-7 (1999).

21. McCracken, S. et al. Diversity in homeostatic calcium set points predicts retinal ganglion cell survival following optic nerve injury in vivo. Cell Reports 42, 113165, DOI: 10.1016/j.celrep.2023.113165 (2023).

22. Pietrucha-Dutczak, M., Amadio, M., Govoni, S., Lewin-Kowalik, J. & Smedowski, A. The role of endogenous neuroprotective mechanisms in the prevention of retinal ganglion cells degeneration. Front. Neurosci. 12, 834, DOI: 10.3389/fnins.2018.00834 (2018).

23. Kasatkina, L. A. & Verkhusha, V. V. Transgenic mice encoding modern imaging probes: Properties and applications. Cell Reports 39, DOI: 10.1016/j.celrep.2022.110901 (2022).

24. Nejat, H., Sherfey, J. & Bastos, A. M. Predictive routing emerges from self supervised stochastic neural plasticity. bioRxiv 2024.12.31.630823, DOI: 10.1101/2024.12.31.630823 (2024).

25. Sekirnjak, C. et al. Electrical stimulation of mammalian retinal ganglion cells with multielectrode arrays. J. Neurophysiol. 95, 3311–3327, DOI: 10.1152/jn.01168.2005 (2006).

26. Wong, R. C. S., Cloherty, S. L., Ibbotson, M. R. & O’Brien, B. J. Intrinsic physiological properties of rat retinal ganglion cells with a comparative analysis. J. Neurophysiol. 108, 2008–2023, DOI: 10.1152/jn.01091.2011 (2012).

27. Freeman, D. K., Rizzo, J. F. & Fried, S. I. Encoding visual information in retinal ganglion cells with prosthetic stimulation. J. Neural Eng. 8, 035005, DOI: 10.1088/1741-2560/8/3/035005 (2011).

28. Crown, L. M. et al. Theta frequency medial septal nucleus deep brain stimulation increases neurovascular activity in mk-801-treated mice. Front. Neurosci. 18, 1372315, DOI: 10.3389/fnins.2024.1372315 (2024).

29. Dowlatabad, H. M. et al. High-frequency (30 mhz–6 ghz) breast tissue characterization stabilized by suction force for intraoperative tumor margin assessment. Diagnostics 13, 179, DOI: 10.3390/diagnostics13020179 (2023).

30. Householder, N. et al. In-vivo two-photon microscopy in thy1-gcamp6f transgenic rats demonstrates that electric field stimulation directly modulates retinal ganglion cell activity. Investig. Ophthalmol. & Vis. Sci. 66, 4775–4775 (2025).

31. Pahlavan, P. et al. Towards non-invasive electrical stimulation for guided optic nerve regeneration. IEEE Transactions on Neural Syst. Rehabil. Eng. DOI: 10.1109/tnsre.2025.3603560 (2025). Please insert final volume/pages/DOI from the publisher record if already assigned.

32. Simonyan, A. et al. Pulsed asymmetric biphasic electrical fields accelerate peripheral nerve regeneration into split thickness, skin graft donor sites. Plast. Reconstr. Surg. 10–1097, DOI: 10.1097/prs.0000000000012394 (2021).

33. Hadjinicolaou, A. E., Cloherty, S. L., Hung, Y.-S., Kameneva, T. & Ibbotson, M. R. Frequency responses of rat retinal ganglion cells. PLoS One 11, e0157676, DOI: 10.1371/journal.pone.0157676 (2016).

34. Guo, T. et al. Mediating retinal ganglion cell spike rates using high-frequency electrical stimulation. Front. Neurosci. 13, 413, DOI: 10.3389/fnins.2019.00413 (2019).

35. Kim, T. et al. Electric field stimulation directs target-specific axon regeneration and partial restoration of vision after optic nerve crush injury. PLoS One 20, e0315562, DOI: 10.1371/journal.pone.0315562 (2025).

36. Upschulte, E., Harmeling, S., Amunts, K. & Dickscheid, T. Contour proposal networks for biomedical instance segmentation. Med. Image Analysis 77, 102371, DOI: 10.1016/j.media.2022.102371 (2022).

37. Alamalhoda, M., Firoozi, A., Venturino, A. & Siegert, S. traice3d: A prompt-driven transformer based for semantic segmentation of microglial cells from large-scale 3d microscopy images (2025). 10.48550/arXiv.2507.22635.

38. Mehradfar, A. et al. Lantern: A machine learning framework for lipid nanoparticle transfection efficiency prediction (2025). 10.48550/arXiv.2507.03209.

39. Uiterkamp, F. E. S. et al. Optic nerve crush does not induce retinal ganglion cell loss in the contralateral eye. Investig. Ophthalmol. & Vis. Sci. 66, 49–49, DOI: 10.1167/iovs.66.3.49 (2025).

40. Cela, C. J., Lee, R. C. & Lazzi, G. Modeling cellular lysis in skeletal muscle due to electric shock. IEEE Transactions on Biomed. Eng. 58, 1286–1293, DOI: 10.1109/tbme.2010.2103362 (2011).

41. Loizos, K. et al. A multi-scale computational model for the study of retinal prosthetic stimulation. In 2014 36th Annual International Conference of the IEEE Engineering in Medicine and Biology Society, 6100–6103, DOI: 10.1109/embc.2014.6945021 (IEEE, 2014).

42. Loizos, K. et al. Increasing electrical stimulation efficacy in degenerated retina: stimulus waveform design in a multiscale computational model. IEEE Transactions on Neural Syst. Rehabil. Eng. 26, 1111–1120, DOI: 10.1109/tnsre.2018.2832055 (2018).

43. Iseri, E., Kosta, P., Paknahad, J., Bouteiller, J.-M. C. & Lazzi, G. A computational model simulates light-evoked responses in the retinal cone pathway. In 2021 43rd Annual International Conference of the IEEE Engineering in Medicine & Biology Society (EMBC), DOI: 10.1109/embc46164.2021.9630642 (IEEE, 2021).

44. Farzad, S. et al. Impact of retinal degeneration on response of on and off cone bipolar cells to electrical stimulation. IEEE Transactions on Neural Syst. Rehabil. Eng. 31, 2424–2437, DOI: 10.1109/tnsre.2023.3276431 (2023).

45. Paknahad, J., Loizos, K., Humayun, M. S. & Lazzi, G. Responsiveness of retinal ganglion cells through frequency modulation of electrical stimulation: A computational modeling study. In 2020 42nd Annual International Conference of the IEEE Engineering in Medicine & Biology Society (EMBC), 3393–3398, DOI: 10.1109/embc44109.2020.9176125 (IEEE, 2020).

46. Ascoli, G. A. Mobilizing the base of neuroscience data: the case of neuronal morphologies. Nat. Rev. Neurosci. 7, 318–324, DOI: 10.1038/nrn1885 (2006).

47. Fohlmeister, J. F. & Miller, R. F. Impulse encoding mechanisms of ganglion cells in the tiger salamander retina. J. Neurophysiol. 78, 1935–1947, DOI: 10.1152/jn.1997.78.4.1935 (1997).

48. Fohlmeister, J. F., Cohen, E. D. & Newman, E. A. Mechanisms and distribution of ion channels in retinal ganglion cells: using temperature as an independent variable. J. Neurophysiol. 103, 1357–1374, DOI: 10.1152/jn.00123.2009 (2010).

49. Kameneva, T., Meffin, H. & Burkitt, A. N. Modelling intrinsic electrophysiological properties of on and off retinal ganglion cells. J. Comput. Neurosci. 31, 547–561, DOI: 10.1007/s10827-011-0322-3 (2011).

50. Van Welie, I., Remme, M. W. H., Van Hooft, J. A. & Wadman, W. J. Different levels of ih determine distinct temporal integration in bursting and regular-spiking neurons in rat subiculum. The J. Physiol. 576, 203–214, DOI: 10.1113/jphysiol.2006.113944 (2006).

51. Wang, X.-J., Rinzel, J. & Rogawski, M. A. A model of the t-type calcium current and the low-threshold spike in thalamic neurons. J. Neurophysiol. 66, 839–850, DOI: 10.1152/jn.1991.66.3.839 (1991).

52. Chen, T.-W. et al. Ultrasensitive fluorescent proteins for imaging neuronal activity. Nature 499, 295–300, DOI: 10.1038/nature12354 (2013).

53. Dana, H. et al. High-performance calcium sensors for imaging activity in neuronal populations and microcompartments. Nat. Methods 16, 649–657, DOI: 10.1038/s41592-019-0435-6 (2019).

54. Helassa, N., Podor, B., Fine, A. & Török, K. Design and mechanistic insight into ultrafast calcium indicators for monitoring intracellular calcium dynamics. Sci. Reports 6, 38276, DOI: 10.1038/srep38276 (2016).

55. Sanes, J. R. & Masland, R. H. The types of retinal ganglion cells: Current status and implications for neuronal classification. Annu. Rev. Neurosci. 38, 221–246, DOI: 10.1146/annurev-neuro-071714-034120 (2015).

56. Qin, W. et al. Single-compartment models of retinal ganglion cells with different electrophysiologies. Network: Comput. Neural Syst. 29, 74–93, DOI: 10.1080/0954898X.2018.1455993 (2018).

57. Paknahad, J., Loizos, K., Yue, L., Humayun, M. S. & Lazzi, G. Color and cellular selectivity of retinal ganglion cell subtypes through frequency modulation of electrical stimulation. Sci. Reports 11, 5177, DOI: 10.1038/s41598-021-84437-w (2021).

58. Werginz, P., Raghuram, V. & Fried, S. I. The relationship between morphological properties and thresholds to extracellular electric stimulation in α rgcs. J. Neural Eng. 17, 045015, DOI: 10.1088/1741-2552/abab47 (2020).

