## Supplement for "Population-Scale Analysis of Frequency-Dependent Calcium Dynamics in Retinal Ganglion Cells Under Electric Field Stimulation"

Table S1 provides a summary of the stimulation waveform parameters used in our experiments. We tested a total of eleven stimulation conditions, comprising one asymmetric charge-balanced (ACB) waveform and ten sinusoidal waveforms with frequencies ranging from 1 Hz to 10 kHz. The ACB waveform was implemented at 50 Hz, with a cathodic phase of 2 ms duration followed by an anodic phase of 8 ms, ensuring charge balance and a 50% duty cycle. For sinusoidal waveforms, each cycle included equal-duration anodic and cathodic phases, starting with anodic phase and no rest period, and the number of applied pulses was adjusted according to frequency to maintain the same total induced charge in the 10 s stimulation window. This selection of waveforms allowed us to study the waveform-specific and frequency-specific aspects of calcium dynamics in the RGC population.

**Supplementary Table S1.** Electrical stimulation waveform parameters. This table summarizes the duration, induced charge per phase, and total number of applied pulses for each stimulation condition used in this study. The 1:4 ACB waveform features a biphasic structure with a short, high-amplitude cathodic phase followed by a longer, lower-amplitude anodic phase, and includes a 10 ms inter-pulse rest period. Sinusoidal waveforms span a broad frequency range (1 Hz to 10 kHz), with the charge per phase decreasing substantially as frequency increases. Charge per phase was calculated by integrating the absolute current over each half-cycle. DC is a 2s monophasic stimulation with 10V amplitude, the same as the peak stimulation amplitude used for sinusoidal waveforms.

| Waveform | Cathodic Duration | Anodic Duration | Rest Duration | Number of applied pulses |
|---|---|---|---|---|
| Sin 1 Hz | 500 ms | 500 ms | 0 | 10 |
| Sin 2 Hz | 250 ms | 250 ms | 0 | 20 |
| Sin 5 Hz | 100 ms | 100 ms | 0 | 50 |
| Sin 10 Hz | 50 ms | 50 ms | 0 | 100 |
| Sin 20 Hz | 25 ms | 25 ms | 0 | 200 |
| Sin 50 Hz | 10 ms | 10 ms | 0 | 500 |
| Sin 100 Hz | 5 ms | 5 ms | 0 | 1k |
| Sin 3 kHz | 166.6 us | 166.6 us | 0 | 30k |
| Sin 5 kHz | 100 us | 100 us | 0 | 50k |
| Sin 10 kHz | 50 us | 50 us | 0 | 100k |
| 1:4 ACB 50 Hz | 2 ms | 8 ms | 10 ms | 500 |
| 2s DC | 2s | 0 | 0 | 1 |

**Supplementary Table S2.** Summary of ion mechanisms used in the RGCF model

| Channel | Full Name | Main Role |
|---|---|---|
| $g_{\mathrm{Na}}$ | Voltage-gated Sodium ($Na^+$) | Initiates action potentials via fast depolarization |
| $g_{\mathrm{K}}$ | Delayed Rectifier Potassium ($K^+$) | Repolarizes the membrane after a spike |
| $g_{\mathrm{K},A}$ | A-type Potassium ($K^+$) | Rapid, transient repolarization; regulates firing frequency |
| $g_{\mathrm{K},Ca}$ | $Ca^{2+}$-activated Potassium ($K^+$) | Enables afterhyperpolarization and spike adaptation |
| $g_{\mathrm{Ca}}$ | High-voltage-gated Calcium ($Ca^{2+}$) | Drives calcium entry for signaling and activates $g_{\mathrm{K},Ca}$ |
| $g_{\mathrm{h}}$ | Hyperpolarization-activated HCN ($Na^+$/$K^+$) | Provides pacemaking current and stabilizes rest |
| $g_{\mathrm{T}}$ | T-type Calcium ($Ca^{2+}$) | Supports burst firing and rebound depolarization |
| $p_{Ca}$ | Calcium Extrusion Pump | Removes intracellular $Ca^{2+}$; restores basal levels |
